# Autonomic challenge uncovers hidden link between diastolic blood pressure and mental imagery vividness

**DOI:** 10.64898/2026.07.12.738020

**Authors:** Yoko Nagai, Angelo Thomas, Lealahni Woulfe, Sam Wray, Timo L Kvamme, Merlin Monzel, Hugo Critchley, Juha Silvanto

**Affiliations:** Brighton & Sussex Medical School, Department of Neuroscience, University of Sussex, UK; Centre for Cognitive and Brain Sciences, University of Macau, China SAR; Department of Experimental Psychology, University of Oxford, UK; Department of Psychology, University of Bonn, Germany; Department of Psychology, Faculty of Health Sciences, University of Macau, China SAR

**Author notes:** Joint senior authors (Hugo Critchley, Juha Silvanto). **Correspondence Author:** Yoko Nagai, Department of Clinical Neuroscience, Brighton and Sussex Medical School, University of Sussex.

**Keywords:** Mental Imagery, Interoception, Cardiovascular function, Aphantasia

## Abstract

Mental imagery is viewed as a fundamental component of human cognition, supporting memory, emotional processing, future simulation, and conscious experience. Although mental imagery vividness has traditionally been attributed to differences in sensory processing, emerging evidence suggests that internally generated images are also shaped by ongoing bodily and interoceptive signals. Here, we investigated whether cardiovascular physiology contributes to individual differences in mental imagery vividness using a virtual reality paradigm in which participants encoded and reconstructed emotional and neutral scenes during a tilt-table-induced autonomic challenge. Acute autonomic changes induced by a tilt table did not alter imagery vividness. However, elevated diastolic blood pressure during the tilt-up condition predicted reduced mental imagery vividness, with the strongest effects observed for disgust and neutral imagery. Our findings provide direct evidence that the vividness of the “mind’s eye” is linked to cardiovascular physiology and support interoceptive accounts of consciousness in which mental imagery emerges from dynamic interactions between neural and bodily systems. More broadly, these results identify cardiovascular function as a potential physiological contributor to altered imagery experiences, including those observed in aphantasia and related clinical conditions.

**Significance statement:** Visual mental imagery has traditionally been explained as a top-down process driven by the brain. We investigated whether bottom-up physiological signals from the cardiovascular system also contribute to imagery vividness. An autonomic challenge revealed a hidden relationship between diastolic blood pressure and the vividness of emotional mental imagery, indicating that ongoing cardiovascular signals influence internally generated visual experiences. These findings identify bottom-up autonomic physiology as a previously unrecognized contributor to mental imagery and broaden current understanding of how top-down and bottom-up processes interact to shape conscious mental imagery experience.

## Introduction

Mental imagery is the capacity to generate vivid, perceptual-like experiences in the absence of external sensory input, supporting memory, emotional processing, planning, and future simulation^1–3^. One of the earliest modern clinical descriptions of acquired imagery loss provided a particularly important clue regarding its possible physiological basis: in 2010, Zeman and colleagues described patient MX, who developed profound loss of visual imagery following a cardiac procedure, despite preserved visual perception and intact visuospatial task performance^4^. MX reported a sudden inability to voluntarily generate mental imagery after the procedure, leading the authors to characterise the phenomenon as “blind imagination.” This report subsequently contributed to the modern conceptualisation and naming of aphantasia^5^, the condition of having little or no voluntary visual imagery, now estimated to affect roughly 3 to 4% of the population^6^. Intriguingly, many individuals with aphantasia still experience visual dreaming, suggesting that voluntary imagery and spontaneous internally generated imagery may rely on partially dissociable mechanisms^7–9^. Yet a fundamental question raised by MX’s case has received surprisingly little attention: why should a cardiac event eliminate visual imagery? Subsequent research on imagery and aphantasia has focused predominantly on visual cortical, attentional, and cognitive explanations, while the possible contribution of cardiovascular and bodily physiology has received comparatively little systematic investigation.

Neuroimaging studies consistently demonstrate overlap between visual perception and imagery-related activity within occipital and temporal cortices, supporting the view that imagery reflects internally generated sensory representations^10–12^. However, accumulating evidence suggests that imagery vividness cannot be fully explained by visual cortical mechanisms alone. Individuals with aphantasia frequently exhibit intact object recognition and perceptual performance despite profoundly diminished subjective imagery^5,13^. Moreover, behavioural and psychophysiological evidence indicates that imagery vividness depends on broader systems involved in bodily awareness and emotional regulation^13,14^. Recent evidence further supports a network-based account of conscious visualisation rather than one based solely on early visual cortex reactivation^15^.

Emerging perspectives from interoceptive neuroscience and embodied cognition provide a framework for understanding these observations. Interoception refers to the sensing and representation of internal bodily states, including cardiovascular, autonomic, and visceral signals^16,17^. Interoceptive processes contribute to emotional awareness, affective processing, self-representation, and conscious experience^18,19^. Cardiovascular signals also influence cognition, modulating attention, emotional salience, sensory awareness, and memory^20,21^.

Recent theoretical and empirical work further suggests that interoception contributes to mental imagery and aphantasia. We previously proposed that interoception and insula-mediated bodily representations contribute to imagery vividness and the phenomenology of aphantasia^22^. Consistent with this framework, we showed that subjective interoceptive awareness predicts imagery vividness^23^ and is associated with autobiographical memory deficits in aphantasia^24^. We also found that core aphantasia and hypophantasia relate differently to mental health outcomes through subjective interoceptive mechanisms^25^, while acquired aphantasia is associated with autonomic and physiological correlates, including early adversity and neurodevelopmental factors^26^. Together, these findings support the view that mental imagery emerges through dynamic interactions between perceptual and bodily systems rather than visual cortical mechanisms alone.

Nevertheless, an important question remains unresolved: do objective cardiovascular signals directly predict imagery vividness? Although previous work has implicated interoceptive awareness in imagery and aphantasia, direct investigation of cardiovascular physiology has remained scarce despite the original clinical observations of Zeman and colleagues following cardiac intervention. To address this question, participants completed an immersive virtual reality (VR) imagery task while cardiovascular activity was monitored during experimentally manipulated autonomic states using a tilt table. Emotional stimuli varied across neutral and affective categories to determine whether physiological influences on imagery differed according to emotional content (Fig. 1).

**Fig. 1.**
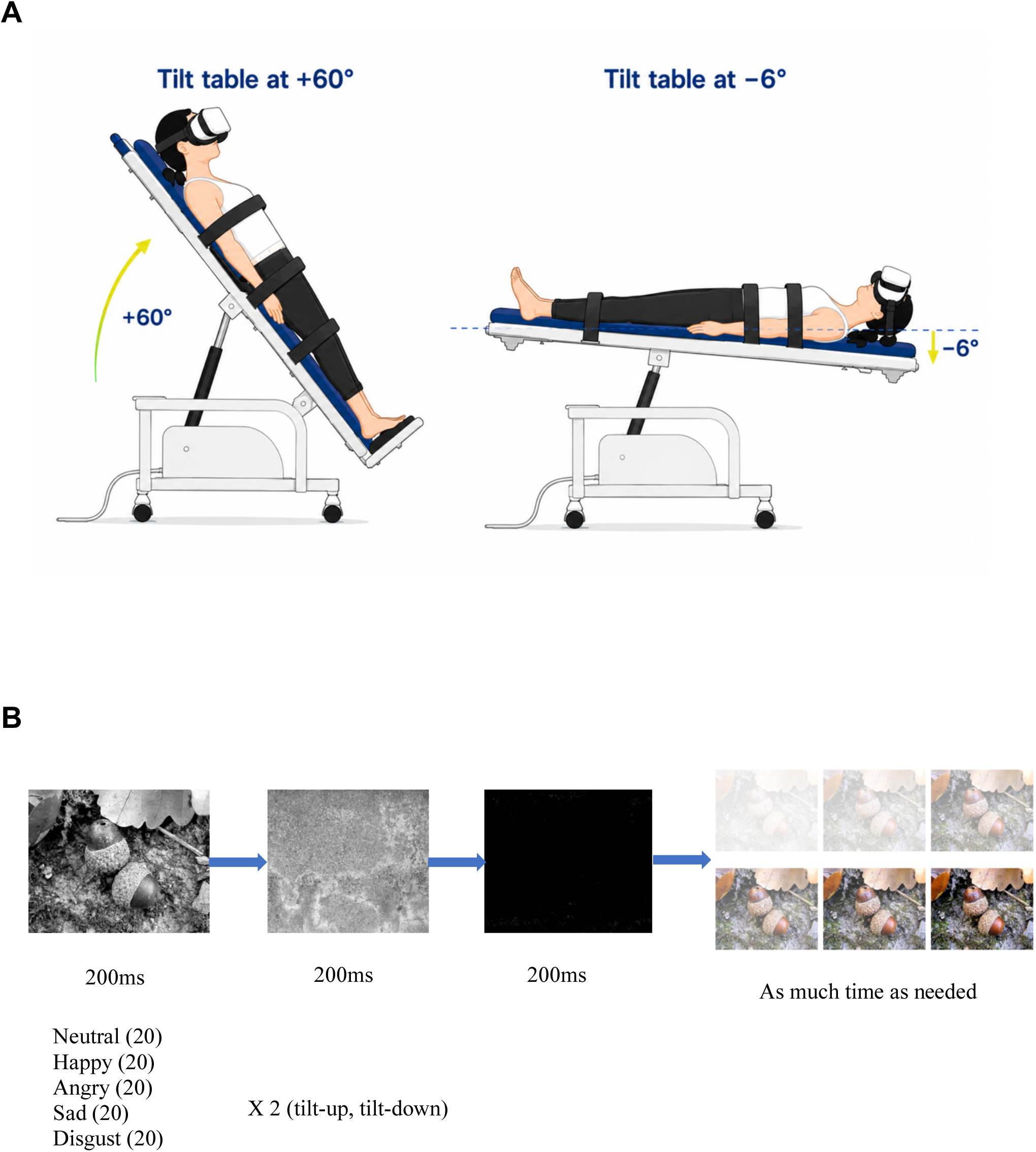
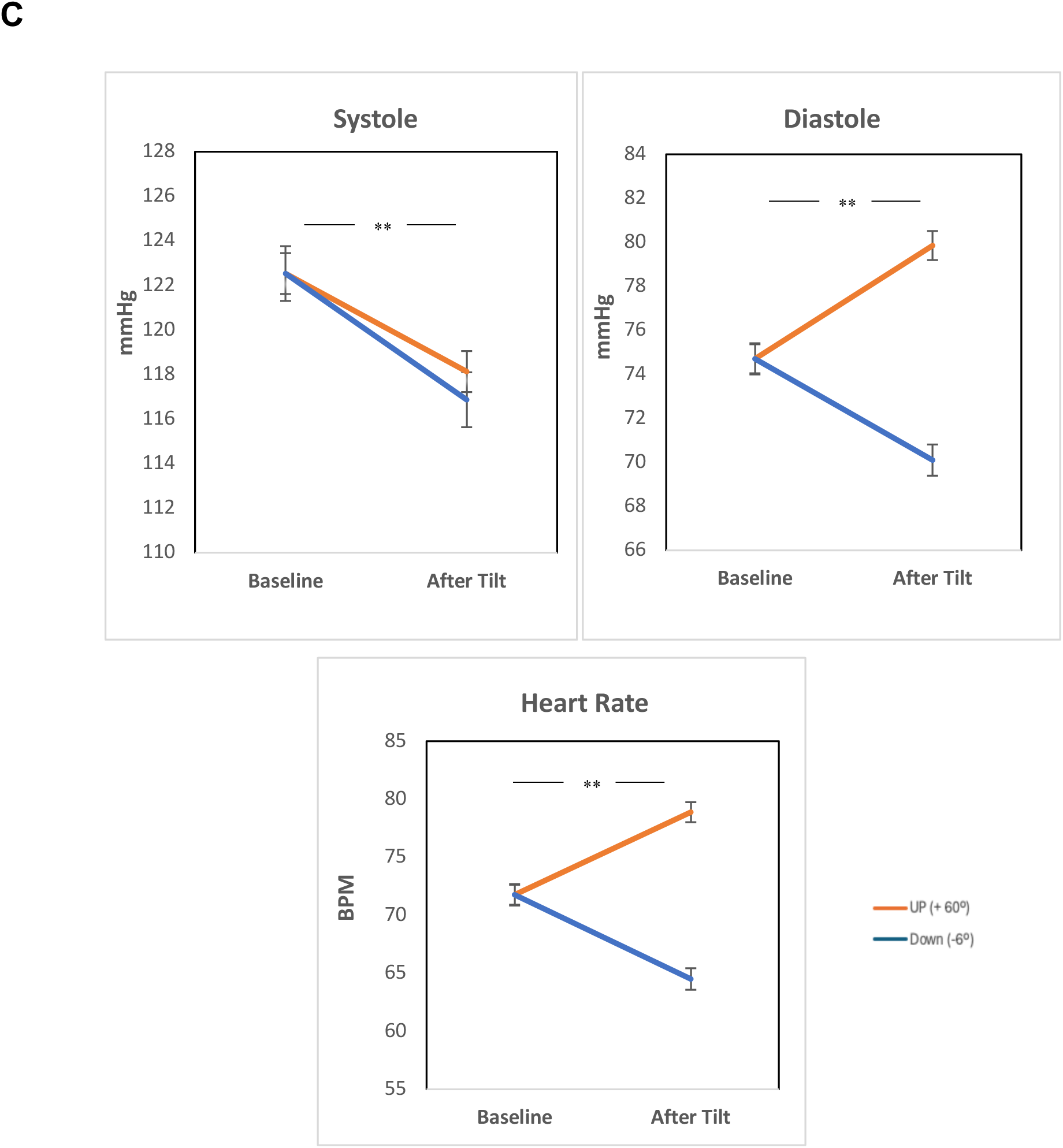
Tilt table and Mental Imagery Vividness Task (MIVT) with VR headset. A) Schematic illustration of the tilt-table positions used in the study. The left panel depicts the tilt-up condition, in which participants were positioned at 60° head-up relative to the horizontal. The right panel depicts the tilt-down condition, in which participants were positioned at 6° head-down (−6°) relative to the horizontal. B) The task involved the presentation of black-and-white images depicting five emotional elements (neutral, happy, angry, sad, and disgusted) for 200 ms. Each image was then followed by a scrambled version of the same image for 200 ms and a blank screen for 200 ms, serving as a priming sequence. Participants were subsequently asked to recall the image in colour and select the corresponding level of vividness from a six-point graded vividness scale. C) Blood pressure and heart rate responses following the imagery task under baseline, −6° head-down tilt, and +60° head-up tilt conditions. Relative to baseline, the −6° head-down tilt was associated with increased systolic and diastolic blood pressure and an elevated heart rate. In contrast, the +60° head-up tilt was associated with changes in blood pressure and a reduced heart rate, reflecting cardiovascular adaptations to altered body position.

We hypothesised that cardiovascular physiology, particularly blood pressure regulation and its putative modulation through baroreceptor–brain interactions, would predict individual differences in imagery vividness. More broadly, we tested whether conscious imagery depends not only on visual cortical mechanisms but also on ongoing physiological state. By combining immersive imagery with physiological monitoring and autonomic modulation, the present study provides one of the first systematic empirical investigations of a question first raised more than a decade ago by the observation of imagery loss following cardiac intervention. The findings support emerging embodied and interoceptive accounts of the “mind’s eye” and suggest that cardiovascular physiology contributes to the phenomenology of conscious mental imagery.

## Results

### Cardiac modulation by tilt table

The tilt-table procedure successfully induced two distinct physiological states. Baseline blood pressure measurements were obtained while participants were seated prior to tilting and were compared with values recorded during the −6° and +60° tilt conditions. Paired-samples t-tests revealed significant differences between baseline blood pressure and both tilt conditions (Fig 1).

Compared to baseline values (systolic: M = 122.53, SD= 14.55; diastolic: M = 74.71, SD= 9.02), the + 60° tilt condition produced significant changes in both systolic blood pressure (M = 118.13, SD= 10.89), *t*(33) = 2.959, *p* = 0.006, and diastolic blood pressure (M = 79.85, SD= 7.88), *t*(33) = −4.785, *p* < 0.001, reflecting the physiological adjustments associated with head-up tilt. Similarly, compared with baseline measurements, the -6° tilt condition resulted in significant and different changes in systolic blood pressure (M = 116.87mmHg, SD=14.50), *t*(33) = 2.941, *p* = 0.006, and diastolic blood pressure (M = 70.10, SD= 8.34), *t*(33) = 3.883, *p* < 0.001, indicating a measurable cardiovascular response to the head-down position.

Heart rate also differed across tilt conditions. Relative to baseline heart rate (M =71.76 bpm, SD= 11.86), heart rate increased significantly during the +60° tilt condition (M = 78.88, SD= 10.38), *t*(33) = −4.580, *p* < 0.001, whereas it decreased during the -6° tilt condition (M =64.51 bpm, SD= 11.00), *t*(33) = 5.556, *p* < 0.001, compared with the baseline). Together, these findings confirm that the tilt-table intervention effectively modified cardiovascular function, resulting in significant blood pressure and heart rate change across the tested positions and indicating the successful induction of an altered autonomic state.

### Task-based and Questionnaire-based imagery vividness

There was a significant correlation between task-based VR Mental Imagery Vividness Task (MIVT) total score (across emotional scenes) and questionnaire-based VVIQ-2 score in both tilt-up (r= 0.687, p = < 0.001) and tilt-down (r= 0.687, p = < 0.001) conditions, indicating that the individuals’ metacognitive appraisal of the vividness of their mental imagery was generally accurate (Fig 2).

**Fig. 2.**
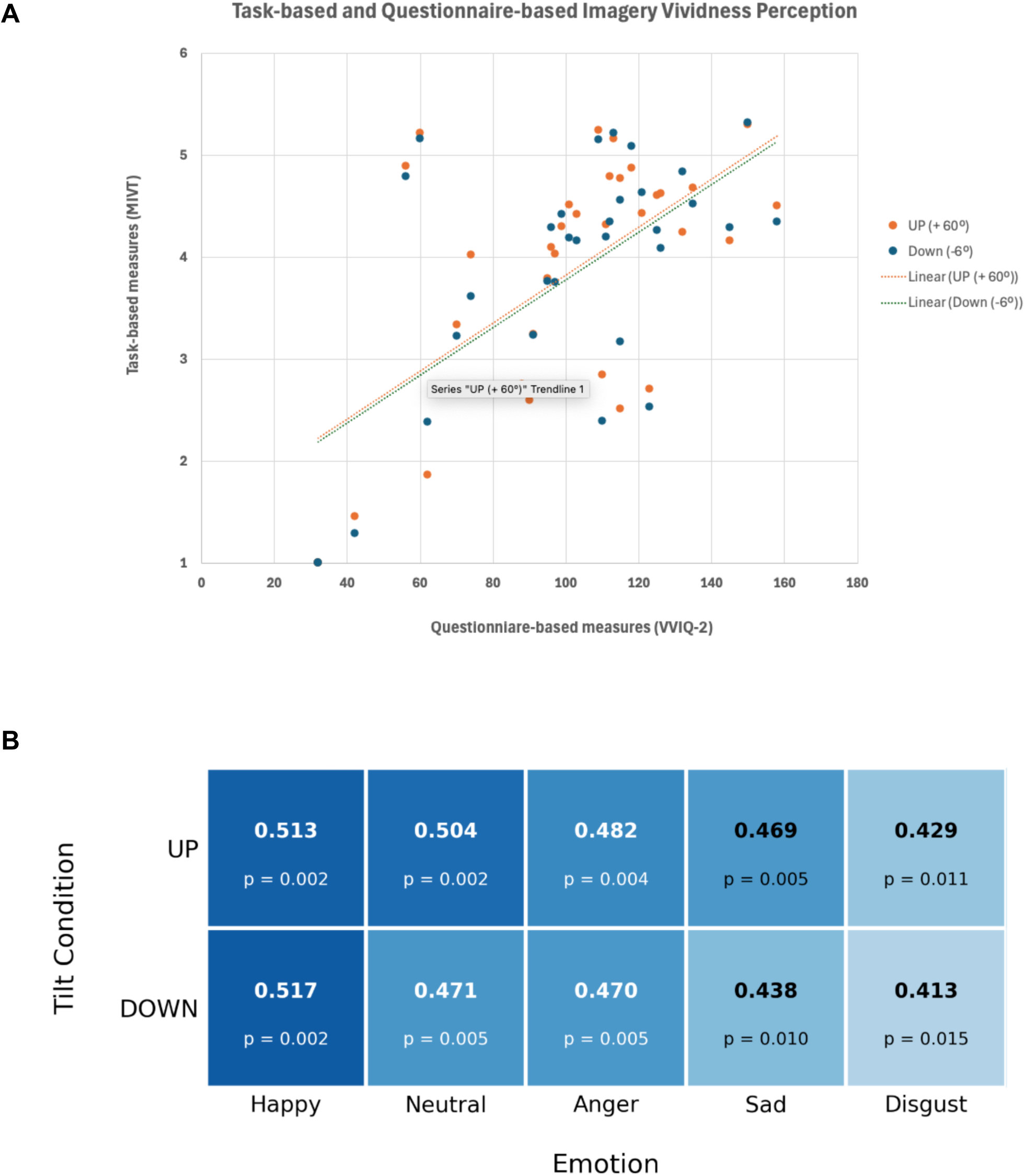
Mental imagery vividness perception. A) Correlation between Task-based VR image recall performance (Mental Imagery Vividness Task: MIVT) and Questionnaire-based mental imagery vividness as measured by the VVIQ-2 for total imagery scores. A significant positive association was observed in both tilt-up and tilt-down conditions. B) Correlations between VVIQ and different emotional categories of happiness, anger, neutral affect, disgust and sadness. Correlation coefficient and p value.

### Effect of emotion and induced acute physiological change on mental imagery vividness

A two-way repeated measures analysis of variance (ANOVA) was performed to assess the effects of emotion (happy, sad, anger, disgust and neutral) and tilt (tilt-up/tilt-down) on average imagery vividness scores.

Although there was no significant main effect of tilt condition (tilt-down and tilt-up) (F(1, 33) =0.98, p= 0.33, η^2^=0.029), we observed a main effect of emotional imagery (happy, sad, anger, disgust and neutral) (F(2.5, 83.3) = 8.26, p<0.001, η^2^=0.20). There was no two-way interaction of Tilt x Emotion (F(2.6, 86.0) = 0.654, p= 0.56, η^2^=0.019).

Post hoc paired *t*-tests revealed that the average imagery vividness score of scenes evoking disgust was lower than the average imagery vividness scores of images evoking happiness (Up: *t*(33) = 3.064, *p* = 0.004, Down: *t*(33) = 3.659, *p* < 0.001), sadness (Up: *t*(33) = 3.233, *p* = 0.003, Down: *t*(33) = 2.659, *p* = 0.012), anger (Up: *t*(33) = 3.070, *p* = 0.004, Down: *t*(33) = 3.593, *p* = 0.001) or neutral affect (Up: *t*(33) = 3.568, *p* = 0.001, Down: *t*(33) = 3.546, *p* < 0.001) in both tilt-up and tilt-down conditions as shown in Fig 3 A.

**Fig. 3.**
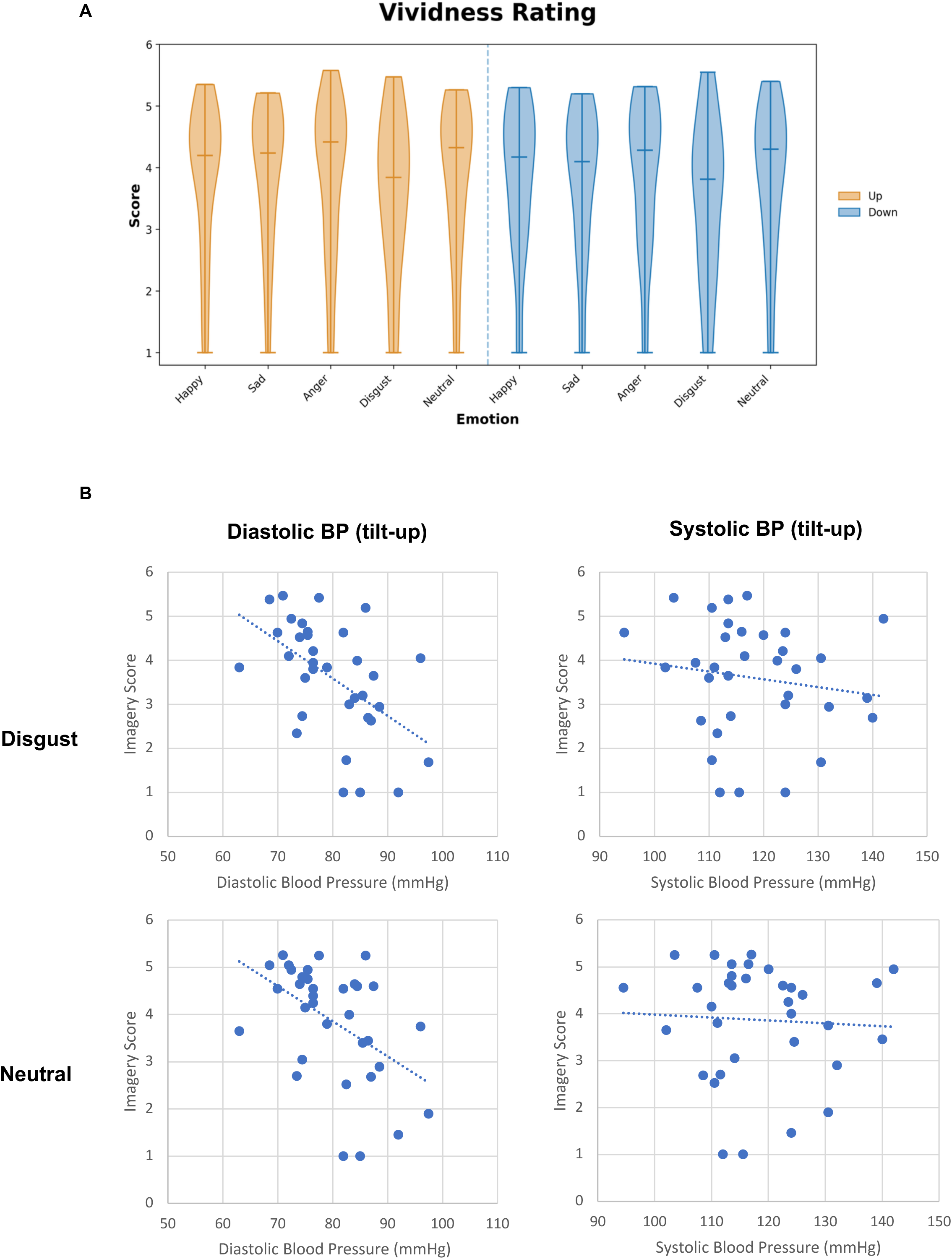
Mental imagery vividness in relation to emotional states and blood pressure. A) Effect of emotion on mental Imagery recall assessed using the Mental Imagery Vividness Task. The violin plot illustrates the mean imagery vividness scores across emotional categories and the distribution of scores within each category in both tilt-up and tilt-down conditions. Images that evoked disgust were associated with significantly lower vividness ratings than images that evoked happiness, anger, neutral affect, or sadness in both conditions. B) Significant associations were found between diastolic blood pressure and disgust (r=- 0.52, p=0.001) and neutral mental imagery (r=-0.47, p=0.005) in tilt-up condition. In contrast, systolic blood pressure showed no significant associations with these imageries. This pattern suggests that the observed effects may be more closely related to changes in vascular resistance and autonomic regulation, which are reflected in diastolic blood pressure, rather than to changes in cardiac contractile activity, which are more strongly associated with systolic blood pressure.

### Link between cardiovascular function, mental imagery vividness and psychological states

A distinct association emerged between objective mental imagery vividness and diastolic blood pressure, observed exclusively during the post–tilt-up condition. Specifically, higher diastolic blood pressure was significantly correlated with reduced imagery vividness during disgust imagery (r = −0.52, p = 0.001) and neutral imagery (r = −0.47, p = 0.005), with both correlation remaining significant following Holm– Bonferroni correction for multiple comparisons (Fig 3 B, Table 1). Similar inverse associations were observed for happy (r = −0.41, p = 0.016), anger (r = −0.37, p = 0.030), and sad imagery (r = −0.35, p = 0.046); however, these relationships did not survive correction for multiple comparisons. Notably, all observed associations were negative in direction, suggesting a broader tendency for elevated diastolic blood pressure to be associated with diminished imagery vividness across emotional imagery conditions.

Supporting the specificity of this finding, only a modest association was observed between imagery vividness and baseline diastolic blood pressure, with neutral imagery showing a negative correlation prior to correction for multiple comparisons in both tilt-up (r = −0.36, p = 0.040) and down (r = −0.39, p = 0.023) conditions. In contrast, no significant relationships were identified between imagery vividness and either systolic blood pressure or heart rate across any emotional condition. Furthermore, the magnitude of evoked changes in blood pressure from baseline to the end of the tilt-up condition was not itself associated with imagery vividness, indicating that the observed effects were not driven by the degree of cardiovascular reactivity per se. Similarly, a heuristic indicator of cardiac output, calculated as pulse pressure × heart rate based on the Windkessel relationship between pulse pressure and stroke volume under assumptions of relatively constant arterial compliance^27^, was not significantly related to imagery vividness, with the exception of a nominal negative correlation for anger imagery (r = −0.37, p = 0.030) that did not survive correction for multiple comparisons. Collectively, these findings suggest a specific and potentially important role for diastolic blood pressure, rather than broader cardiovascular indices, in relation to mental imagery vividness.

Heart beat timing was also recorded during both the tilt-up and tilt-down conditions while participants completed the mental imagery vividness task. Heart rate variability (HRV) metrics were then derived for each experimental condition. Significant differences between tilt-up and tilt-down conditions were observed for HRV indices Root Mean Square of Successive Differences (RMSSD) (t(31) = 2.23, p = 0.03) and high-frequency (HF) FFT power (t(31) = 5.01, p < 0.001), indicating altered autonomic (parasympathetic) cardiac drive between postures. Low-frequency (LF) FFT power did not differ significantly between the tilt-up and tilt-down conditions. Correlation analyses revealed no significant associations between HRV measures (RMSSD, HF FFT, or LF FFT) and imagery vividness ratings across the emotional conditions.

There were no significant correlations between task-based mental imagery vividness and the psychological questionnaire measures. However, before correction for multiple comparisons, the score on the Multidimensional Assessment of Interoceptive Awareness (MAIA) axis of Emotional Awareness was negatively associated with vividness ratings of happy (r = −0.36, p = 0.04) and disgust (r = −0.37, p = 0.03) emotional scenes in the tilt-up condition, and for disgust emotion in the tilt-down condition (r = −0.34, p = 0.05). In addition, vividness ratings for disgust emotion in the tilt-down condition were negatively associated with the MAIA Not-Worrying subscale (r = −0.37, p = 0.03).

## Discussion

The present study investigated whether indices of cardiovascular physiology contribute to the vividness of emotional mental imagery. We demonstrated a significant bottom-up influence of autonomic physiology on imagery vividness, with diastolic blood pressure predicting imagery performance under experimentally manipulated autonomic conditions (tilt-up). In addition, we found that emotional content influenced imagery vividness, with disgust-related scenes producing significantly lower vividness ratings than other emotional categories. Together, these findings suggest that mental imagery is shaped not only by top-down cognitive and perceptual mechanisms but also by ongoing physiological signals arising from the body (bottom-up, interoceptive signals).

The influence of autonomic activity on mental imagery has traditionally been viewed as a top-down phenomenon. A substantial literature shows that mentally simulating emotional experiences, particularly fear- and anxiety-provoking scenarios, elicits autonomic responses including changes in heart rate, blood pressure, respiration, skin conductance, and peripheral vascular activity^28–30^. Mental imagery has therefore been described as a simulation of reality that activates physiological systems in ways resembling actual perception and experience^30^. Early physiological studies further showed that autonomic response patterns during imagery closely resemble those observed during corresponding real-world actions^28^. Together, these findings indicate that imagery exerts powerful effects on autonomic regulation.

The present findings extend this literature by demonstrating the converse relationship. Rather than autonomic activity simply resulting from imagery, cardiovascular physiology itself predicted imagery vividness. This effect was specific to diastolic blood pressure during sympathetic activation induced by the tilt-up condition. Tilt-up causes blood pooling in the lower body, requiring compensatory cardiovascular adjustments that increase baroreceptor engagement. The present findings therefore suggest that cardiovascular afferent signalling contributes directly to the subjective vividness of conscious imagery. A potential explanation for this relationship lies in the physiology of blood pressure regulation. Arterial baroreceptors in the carotid sinus and aortic arch continuously monitor arterial pressure and relay afferent signals via the nucleus tractus solitarius to forebrain regions involved in predictive allostasis and autonomic regulation^31^. These pathways include the insular cortex, anterior cingulate cortex, and amygdala, which are increasingly recognised as key regions for interoception, emotional awareness, self-representation, and conscious experience. Through these ascending pathways, cardiovascular information is integrated with ongoing sensory and mnemonic processing^16,31^. Consequently, fluctuations in blood pressure may influence conscious experience by altering the physiological information available to higher-order cognitive systems.

The specificity of the effect to diastolic blood pressure may provide further insight into the underlying mechanism. Unlike systolic blood pressure, which primarily reflects cardiac output and ventricular contraction, diastolic pressure is more strongly influenced by peripheral vascular resistance and vascular tone^32^, providing a relatively stable index of cardiovascular regulation and baroreceptor engagement. Variation in peripheral vascular resistance may therefore influence the strength of cardiovascular afferent signalling reaching cortical networks involved in imagery generation. Alternatively, individuals with higher diastolic blood pressure during orthostatic challenge may represent a distinct autonomic phenotype characterised by greater peripheral vascular resistance and altered cardiovascular–brain interactions. Cardiovascular afferent signals arising from arterial baroreceptors influence central nervous system function beyond blood pressure regulation, contributing to sensory processing, attention, and interoceptive awareness through projections to the central autonomic network^16,31^. Evidence that baroreceptor activation can attenuate pain perception and modulate sensory processing suggests that cardiovascular afferent signalling may exert broader inhibitory influences on cortical function^33,34^, raising the possibility that individual differences in cardiovascular signalling also influence the neural processes supporting internally generated imagery representations. The precision and richness of these representations may therefore depend on the quality and continuity of ascending cardiovascular signals.

Importantly, imagery vividness was predicted not by the change in diastolic blood pressure from baseline to tilt-up, but by the absolute diastolic blood pressure maintained during the task, indicating that the effect reflects sustained peripheral vascular resistance during imagery generation rather than cardiovascular reactivity to orthostatic challenge. Thus, orthostatic challenge unmasked a latent association between diastolic blood pressure and imagery vividness, suggesting that imagery depends more on sustained cardiovascular state than moment-to-moment cardiovascular fluctuations. Baseline diastolic blood pressure was also correlated with neutral imagery vividness in both tilt conditions, suggesting a role in basic imagery generation. Upright autonomic challenge may therefore increase the dependence of mental imagery on cardiovascular regulation. However, this interpretation remains speculative given the correlational nature of the data, and future studies that directly manipulate baroreceptor input during imagery are needed to test this hypothesis. This distinction may also explain why heart rate (HR) and heart rate variability (HRV) did not predict imagery despite differing significantly between tilt conditions. Under orthostatic sympathetic activation, HR and HRV primarily reflect beat-to-beat cardiac timing, whereas blood pressure more directly reflects the sustained vascular regulation associated with baroreceptor signalling. Consequently, HR and HRV may be less sensitive markers of the interoceptive signals relevant to imagery generation.

A lack of cerebral perfusion is unlikely to explain the findings. Elevated diastolic blood pressure, but not systolic blood pressure, predicted imagery vividness, despite systolic pressure being more closely related to cerebral perfusion. Moreover, imagery vividness was not enhanced during head-down tilt, which would be expected if increased cerebral perfusion improved imagery.

A second major finding was that disgust-related scenes produced significantly lower imagery vividness ratings than other emotional categories. Although emotional imagery generally enhances subjective experience, disgust is uniquely associated with contamination avoidance, visceral sensations, and behavioural withdrawal^35^. Rather than promoting detailed visual elaboration, disgust may favour rapid categorisation and avoidance. This interpretation is consistent with the role of the insular cortex in disgust perception and interoception^36,37^, suggesting that disgust imagery preferentially engages visceral rather than visual-perceptual representations, thereby reducing imagery vividness.

In conclusion, the present study provides the first systematic investigation of the relationship between cardiovascular physiology and emotional mental imagery vividness using an immersive virtual reality paradigm with continuous physiological monitoring. By manipulating autonomic state while minimising environmental differences between tilt conditions, we demonstrate that diastolic blood pressure predicts imagery vividness under conditions of increased baroreceptor engagement and that disgust-related emotional content reduces imagery vividness. Together, these results suggest that conscious imagery is shaped not only by visual cortical and cognitive mechanisms but also by ongoing cardiovascular physiology. More broadly, the findings support emerging embodied accounts of the mind’s eye and highlight cardiovascular signalling as a previously underexplored contributor to conscious imagery.

Several limitations should be acknowledged. First, the observed associations are correlational and do not establish causal mechanisms linking blood pressure regulation to imagery vividness. Second, it remains unclear whether these effects extend beyond emotional imagery. Third, although blood pressure provided a useful physiological marker, future work incorporating measures of baroreceptor sensitivity, heart rate variability, cerebral blood flow, and neuroimaging would help clarify the neural mechanisms underlying these effects.

## Materials and Methods

### Participants

Thirty-five adults participated in the mental imagery study. However, one participant was excluded from the analysis due to a technical fault during data collection, leaving a final sample of 34 participants (M = 26.37 years, SD = 8.95). Data from three participants were excluded from the HRV analysis because of recording artefacts. 17 identified as men, 16 as women and 1 as non-binary. They were recruited via personal contact and posters across the university campus and all met the inclusion criteria. The inclusion criteria required for participation includes the following: being age 18-70; being fluent in English; having no known cardiovascular or neurological conditions; and not currently taking medications that affect heart rate or blood pressure. All participants provided written informed consent prior to participation. The study was conducted in accordance with the ethical principles of the Declaration of Helsinki, and ethical approval was obtained the Brighton and Sussex Medical School Research Governance and Ethics Committee (ER/BSMS9RN9/1) before the commencement of the study. Participants were compensated £30 for their time.

### Questionnaires

Before conducting the main study, participants were asked to fill in the following questionnaires.

The State-Trait Anxiety Inventory – State portion (STAI-S)^38^ is a validated 20-item self-report questionnaire that uses a 4-point Likert scale to measure a person’s temporary, immediate, and situational feelings of apprehension, tension, and nervousness at a specific moment in time.

The Positive and Negative Affect Schedule (PANAS)^39^ is a 20-item self-report questionnaire that uses a 5-point Likert scale to measure a person’s positive and negative affects.

The Vividness of Visual Imagery Questionnaire-2^40^ is a 32-item self-report questionnaire that uses a 5-point Likert scale to measure the clarity and intensity of a person’s mental images. VVIQ-2 was performed to see how the individuals perceive their mental imagery vividness.

The Multidimensional Assessment of Interoceptive Awareness-2 (MAIA-2)^41^ is a 37- point self-report questionnaire that uses a 6-point Likert scale to measure subjective interoception.

Toronto Alexithymia Scale-20 (TAS-20)^42^ is a 20-item self-report questionnaire that uses a 5-point Likert scale to identify and measure alexithymia.

### Mental Imagery Vividness Task and Procedure

The task was conducted using a virtual reality (VR) headset (Meta Quest 3S) to present instructions and visual stimuli across different body positions. VR has been used extensively in psychological and neuroscience research to provide standardized visual presentation and experimental control50,51. Participants performed the task on a tilt table, enabling controlled manipulation of body orientation and cardiovascular function (see below). A computer display was streamed to the VR headset using Virtual Desktop software. The experiment was programmed using PsychoPy and Spyder (V 6.1.3).

During each trial, participants viewed a greyscale target image (200 ms), a greyscale scrambled image (200 ms), and a black screen (200 ms), before imagining the target image in its original colours. They then selected, without time limit, which of six colour-opacity levels (2 × 3 response grid; Figure 1) best matched the vividness of their mental image. The greyscale target image and scrambled image minimised short-term visual memory by reducing the contribution of colour information^43^ and introducing visual interference^44^. A total of 200 images were selected from the International Affective Picture System (IAPS) and the Open Affective Standardized Image Set (OASIS), equally distributed across five emotional categories (happiness, anger, sadness, disgust, and neutrality).

Participants completed 100 trials in each experimental condition (tilt-up and tilt-down; see below), with images presented in random order. Participants were randomly assigned to complete either the tilt-up or tilt-down condition first. After a five-minute rest period with the VR headset removed, they completed the remaining condition. Condition order was counterbalanced across participants to minimise order effects. Testing was conducted in the same laboratory under controlled environmental conditions, with the lights off, blinds closed, and windows shut throughout data collection.

### Tilt table

The imagery task was conducted on a tilt table, a well-established, safe, and non-invasive method for eliciting controlled cardiovascular and autonomic responses through postural manipulation. Head-up tilt testing is widely used to examine orthostatic cardiovascular regulation because changes in body position produce predictable alterations in blood pressure, heart rate, venous return, and autonomic activity^45^. Blood pressure responses are widely used as physiological markers of autonomic state. Accordingly, the present study assumed that postural manipulation would induce sufficient changes in blood pressure to alter autonomic state. Two tilt-table positions were used: standing and lying. In the standing condition (tilt-up), participants performed the imagery task at a 60° head-up tilt. In the lying condition (tilt-down), participants performed the task at a 6° head-down tilt (Figure 1).

### Cardiac monitoring (blood pressure and ECG)

Blood pressure was measured using an automatic monitor (Omron M2 Basic HEM- 7121J-E) at baseline (seated before the imagery task), during the tilt-up condition, and during the tilt-down condition after completion of half of the trials in each condition. Electrocardiogram (ECG) activity was recorded continuously using a CED 9201 data acquisition interface and Spike software (Cambridge Electronic Design, UK). ECG signals were processed using R-DECO (MATLAB) for R-peak detection and NeuroKit2 (Python) for heart rhythm and heart rate variability (HRV) analyses. RR intervals were analysed using standard HRV procedures. Artefacts were identified automatically and verified by visual inspection. Time-domain HRV was quantified as the root mean square of successive differences (RMSSD), and frequency-domain HRV was assessed using fast Fourier transform (FFT) spectral analysis. All analyses were performed on artefact-corrected RR interval data.

### Statistical analysis

A paired-samples t-test was used to examine differences in baseline blood pressure and other cardiac parameters between the two tilt-table conditions (−6° and +60°). For the imagery task, a two-way repeated-measures analysis of variance (ANOVA) was conducted to assess within-subject differences in imagery vividness across physiological states induced by the tilt-table conditions and across five emotion categories. The factors included two tilt-table conditions (tilt-up and tilt-down) and five emotions (happiness, sadness, anger, disgust and neutral), allowing for the assessment of main effects and interaction effects. Greenhouse–Geisser corrections were applied to account for violations of sphericity. Post hoc analyses were performed using paired-samples t-tests to investigate significant within-subject differences. Correlations between variables were assessed using Spearman’s rank correlation coefficient, as at least one variable did not satisfy the assumption of normality. Holm–Bonferroni correction was used for multiple comparison at the cut of at P = 0.0167.

## Acknowledgement

This work was supported by the Internal Research Fund of Brighton and Sussex Medical School, University of Sussex. The authors thank Dr. Vasso Anagnostopoulou for her statistical advice and Joel Patcheitt for his valuable technical support.

